# Systemic and mucosal antibody secretion specific to SARS-CoV-2 during mild versus severe COVID-19

**DOI:** 10.1101/2020.05.21.108308

**Authors:** Carlo Cervia, Jakob Nilsson, Yves Zurbuchen, Alan Valaperti, Jens Schreiner, Aline Wolfensberger, Miro E. Raeber, Sarah Adamo, Marc Emmenegger, Sara Hasler, Philipp P. Bosshard, Elena De Cecco, Esther Bächli, Alain Rudiger, Melina Stüssi-Helbling, Lars C. Huber, Annelies S. Zinkernagel, Dominik J. Schaer, Adriano Aguzzi, Ulrike Held, Elsbeth Probst-Müller, Silvana K. Rampini, Onur Boyman

## Abstract

**Background:** Infection with the severe acute respiratory syndrome coronavirus 2 (SARS-CoV-2) causes an acute illness termed coronavirus disease 2019 (COVID-19). Humoral immune responses likely play an important role in containing SARS-CoV-2, however, the determinants of SARS-CoV-2-specific antibody responses are unclear.

**Methods:** Using immunoassays specific for the SARS-CoV-2 spike protein, we determined SARS-CoV-2-specific immunoglobulin A (IgA) and immunoglobulin G (IgG) in sera and mucosal fluids of two cohorts, including patients with quantitative reverse-transcriptase polymerase chain reaction (RT-qPCR)-confirmed SARS-CoV-2 infection (n = 56; median age 61 years) with mild versus severe COVID-19, and SARS-CoV-2-exposed healthcare workers (n = 109; median age 36 years) with or without symptoms and tested negative or positive by RT-qPCR.

**Findings:** On average, SARS-CoV-2-specific serum IgA titers in mild COVID-19 cases became positive eight days after symptom onset and were often transient, whereas serum IgG levels remained negative or reached positive values 9–10 days after symptom onset. Conversely, patients with severe COVID-19 showed a highly significant increase of SARS-CoV-2-specific serum IgA and IgG titers as a function of duration since symptom onset, independent of patient age and comorbidities. Very high levels of SARS-CoV-2-specific serum IgA correlated with severe acute respiratory distress syndrome (ARDS). Interestingly, some of the SARS-CoV-2-exposed healthcare workers with negative SARS-CoV-2-specific IgA and IgG serum titers had detectable SARS-CoV-2-specific IgA antibodies in their nasal fluids and tears. Moreover, SARS-CoV-2-specific IgA levels in nasal fluids of these healthcare workers were inversely correlated with patient age.

**Interpretation:** These data show that systemic IgA and IgG production against SARS-CoV-2 develops mainly in severe COVID-19, with very high IgA levels seen in patients with severe ARDS, whereas mild disease may be associated with transient serum titers of SARS-CoV-2-specific antibodies but stimulate mucosal SARS-CoV-2-specific IgA secretion. The findings suggest four grades of antibody responses dependent on COVID-19 severity.

## Introduction

Severe acute respiratory syndrome coronavirus 2 (SARS-CoV-2), the causative agent of coronavirus disease 2019 (COVID-19), is a betacoronavirus related to severe acute respiratory syndrome coronavirus (SARS-CoV) and Middle East respiratory syndrome coronavirus (MERS-CoV).^1–4^ The zoonotic introduction of MERS-CoV and SARS-CoV into the human population resulted in limited outbreaks, whereas the appearance of SARS-CoV-2 has led to a rapidly spreading pandemic. As of May 21, 2020, COVID-19 had affected about 5 million confirmed cases worldwide in more than 213 countries and caused an estimated 330,000 deaths. Several characteristics of SARS-CoV-2 have likely contributed to its rapid spread. These include the ability of SARS-CoV-2 to efficiently replicate in the upper respiratory tract mucosa of humans,^5^ its variable incubation time of about 3–14 days, and the presence of many asymptomatic and presymptomatic SARS-CoV-2-infected individuals producing sufficient amounts of virus for human-to-human transmission.^6–8^ Thus, SARS-CoV-2 infection is frequently unrecognized.

When symptomatic, COVID-19 can range from a mild flu-like illness in about 81% to a severe and critical disease in about 14% and 5% of affected patients, respectively.^9,10^ Mild COVID-19 is characterized by fatigue, fever, sore throat, cough, and mild pneumonia. Severe disease features dyspnea, hypoxia and radiographic evidence of lung involvement of 50% and more. And critical COVID-19 results in acute respiratory distress syndrome (ARDS) with respiratory failure, multiorgan dysfunction, and shock. The World Health Organization (WHO) proposed a classification of symptomatic COVID-19 into (i) mild illness, (ii) mild pneumonia, (iii) severe pneumonia, (iv) ARDS (based on the Berlin definition of ARDS)^11^, and (v) sepsis and septic shock.^12^

Human angiotensin-converting enzyme 2 (ACE2) serves as a cell entry receptor for SARS-CoV-2. Pneumocytes and other host cells expressing ACE2 are therefore particularly susceptible to infection by SARS-CoV-2. Mechanistically, SARS-CoV-2 binds to ACE2 via the receptor-binding domain (RBD) of the S1 subunit of its surface spike (S) glycoprotein.^3,13^ Thus, humoral immunity targeting the S protein could interfere with SARS-CoV-2 infection, as evidenced from serological studies.^14,15^

As with other coronaviruses, symptomatic SARS-CoV-2 disease causes an acute infection with activation of the innate and adaptive immune systems. The former leads to the release of several pro-inflammatory cytokines, including interleukin-6, whereas other anti-viral cytokines, such as the type I and III interferon pathways, are hampered by coronaviruses, including SARS-CoV and SARS-CoV-2.^16–18^ Subsequently, B and T cells become activated, resulting in the production of SARS-CoV-2-specific antibodies, comprising immunoglobulin M (IgM), immunoglobulin A (IgA), and immunoglobulin G (IgG). Whereas coronavirus-specific IgM production is transient and leads to isotype switch to IgA and IgG, these latter antibody subtypes can persist for extended periods in the serum and in nasal fluids.^19^ Whether SARS-CoV-2-specific IgG antibodies correlate with virus control is a matter of intense discussions.^14,15,20^

Unlike the internal nucleocapsid protein (NC) of SARS-CoV-2, that shares about 90% amino acid sequence homology with the NC of SARS-CoV, the S1 subunit shares only 64% amino acid sequence homology and shows limited homology with other human coronaviruses, such as 229E, NL63, OC43, and HKU1, which use different viral entry receptors.^3,21^ Thus, antibodies generated to previous coronavirus infections are unlikely to cross-react with the S1 protein of SARS-CoV-2 and should therefore not significantly account for any seroreactivity toward the S1 subunit.^21^

Despite intensive research efforts, several determinants of SARS-CoV-2-specific antibody production remain ill-defined, such as its relation to COVID-19 severity, disease duration, patient age, and comorbidities. There is also a paucity of knowledge on SARS-CoV-2-specific IgA and IgG antibodies at mucosal sites and how their levels correlate with COVID-19 parameters. And, finally, it is unclear whether tissue-associated IgA and IgG secretion, rather than their systemic production, might be evident in SARS-CoV-2-exposed individuals undergoing mild disease.

## Methods

### Human subjects and patient characteristics

Following written informed consent, patients and healthcare workers were recruited for sampling of blood and mucosal secretions. We studied two cohorts, including patients with quantitative reverse-transcriptase polymerase chain reaction (RT-qPCR)-confirmed SARS-CoV-2 infection (n = 56; median age 61 years) with mild versus severe COVID-19, and healthcare workers possibly exposed to SARS-CoV-2-infected patients (termed HCW cohort; n = 109; median age 36 years) with or without symptoms who tested negative or positive for SARS-CoV-2 by RT-qPCR. Possibly exposed was defined as a contact with a confirmed COVID-19 patient for more than 15 minutes, at less than 2 meters, and without adequate safety measures. Because of pre-existing comorbidities, seven patients were under immunosuppressive treatments, which comprised corticosteroids at stable doses, whereas patients taking B cell-depleting agents, such as rituximab,^22^ were excluded. For longitudinal analyses of serum and mucosal SARS-CoV-2-specific antibody responses two subjects with mild COVID-19 were sampled repeatedly during their disease course. Our COVID-19 patients were classified according to the WHO criteria^12^ into (a) mild cases, comprising mild illness and mild pneumonia, versus (b) severe cases, including severe pneumonia and ARDS; our cohort did not contain any patients with sepsis or septic shock. The study was approved by the Cantonal Ethics Committee of Zurich (BASEC #2016-01440).

### Collection of serum, tears, nasal fluid and saliva

A subgroup of the HCW cohort (termed HCW mucosal subgroup; n = 33) volunteered to be sampled for blood as well as, simultaneously, tears, nasal fluid, and saliva. Venous blood samples were collected in BD Vacutainer CAT serum tubes (Becton Dickinson). Tears were sampled using filter paper produced for Schirmer tear tests (HS Clement Clarke Ophtalmic). Nasal fluids were collected by inserting a dry soft tissue into the nasal cavities for 5 minutes. Unstimulated saliva was collected for 5 minutes.

### IgA and IgG immunoassays

A commercial enzyme-linked immunosorbent assay (ELISA) specific for the S1 protein of SARS-CoV-2 was used according to manufacturer’s instructions (Euroimmun SARS-CoV-2 IgA and IgG immunoassay) and validated by using serum samples of hospitalized patients with confirmed COVID-19 as positive controls and serum samples collected prior to the COVID-19 pandemic as negative controls. The results showed a specificity for anti-S1 IgA of >95% and anti-S1 IgG >99%, which is in accordance with recently published data.^23^ Serum samples were analyzed at a 1:100 dilution and mucosal samples at a 1:5 dilution. For serum IgA, optical density (OD) ratios of 1.1–2.0 were considered borderline positive and values above 2.0 positive. For serum IgG, OD ratios of 0.8–1.1 were considered borderline positive and values above 1.1 positive.

Furthermore, we assessed the samples of the HCW mucosal subgroup using an in-house immunoassay for IgA and IgG against S protein extracellular domain (ECD), RBD, and NC. Mucosal samples were pre-diluted 1:2 in sample buffer (PBS Tween-20 0.1%, 1% milk) and serum was pre-diluted 1:20 in sample buffer and transferred to antigen-coated 1536-well assay plates using acoustic dispensing technology (Labcyte ECHO 555) with serial dilutions ranging from 1:5–1:640 (mucosal samples) and 1:50– 1:6400 (serum). Following a two-hour incubation at room temperature, plates were washed 5 times with PBS Tween-20, 0.1%, and HRP-conjugated antibody (peroxidase AffiniPure goat anti-human IgG, Fcγ fragment specific, Jackson) was added in sample buffer. Plates were washed three times with PBS Tween-20, 0.1%, the chromogenic substrate was added and the reaction was stopped with sulphuric acid after three minutes of incubation. ODs were measured at 450 nm in a multimode plate reader (Perkin Elmer EnVision), followed by fitting with a logistic regression and determination of the inflection point of the sigmoidal curve (-log(EC50)). Negative values were depicted as 0.

### Quantitative reverse-transcriptase polymerase chain reaction (RT-qPCR)

Nasopharyngeal swabs were subjected to RT-qPCR using the TaqMan SARS-CoV-2 Assay Kit v2 (Thermo Fischer), the 2019-nCoV CDC qPCR Probe Assay (2019-nCov CDC EUA Kit; Integrated DNA Technologies, Inc.), or the Roche Cobas SARS-CoV-2 Test CE-IVD (Roche) according to manufacturers’ instructions. The cycle threshold (Ct) values for the different SARS-CoV-2 PCR targets were compounded and reported as averages.

### Statistics

Descriptive statistics for the cohort of patients (stratified by mild versus severe disease) and the HCW cohort are presented as median and interquartile ranges for continuous variables, as well as numbers and percentages of total for categorical variables. For the comparison of location parameters in two independent groups, the Wilcoxon’s rank sum test was used, in a version accounting for ties.^24^ For the comparison of more than two independent groups, the non-parametric Kruskal-Wallis test was used. Multiple linear regression models were used to quantify the association between log-transformed IgA and IgG levels as outcomes as well as a set of pre-defined independent variables. These included disease severity, age, duration of symptoms (days) and patient comorbidities. Generalized additive models were used to evaluate potential non-linear relationships of disease duration with the two outcomes, as described above.

Statistical analysis were performed with R (version 3.6.1) and using the packages “coin” and “mgcv”. Graph-Pad Prism was used for visualizations. P-values were adjusted for multiple testing, using the method proposed by Benjamini-Hochberg.^25^ Adjusted p-values were considered statistically significant if smaller than the significance level of α = 0.05. In the HCW mucosal subgroup evidence was quantified on a continuous scale, and these results were considered exploratory.

## Results

### COVID-19 severity, disease duration, and patient age influence SARS-CoV-2-specific serum IgA and IgG secretion

Serum samples of 56 RT-qPCR-confirmed mild (n = 19) and severe (n = 37) COVID-19 cases (Table 1) were assessed for IgA and IgG antibodies toward the SARS-CoV-2 S1 protein by using highly-specific immunoassays. The mean period between reported symptom onset and serum collection were 16.4 days (median 13 days) in the mild COVID-19 group and 20.9 days (median 16 days) in the severe COVID-19 group of patients, respectively, which was not significantly different between these two groups (p = 0.17; **Supplementary Figure S1A**). In comparison to mild COVID-19, patients with severe disease had on average higher serum titers of S1-specific IgA (p = 0.0006) and IgG (p = 0.0009) (Figure 1A). In mild COVID-19 cases, serum titers of S1-specific IgA increased slightly as a function of disease duration (p = 0.003), as calculated from symptom onset to the time of serum sampling (Figure 1B). Likewise, serum titers of S1-specific IgG increased moderately (p = 0.055) in mild cases (Figure 1B). These antibody responses revealed no significant pattern associated with patient age (p = 0.067 for IgA, and p = 0.18 for IgG) (Figure 1C and **Supplementary Figure S1B**), gender (**Supplementary Figure S1C** and **S1D**), pre-existing comorbidities, including hypertension, diabetes mellitus, heart disease, cerebrovascular disease, lung disease, kidney disease, and malignancy, or immunosuppressive treatment (**Supplementary Figure S2** and Table 2).

**Table 1.** Demographic and clinical characteristics of the patient cohort. Patients were divided in mild versus severe COVID-19 cases. Disease severity was defined according to the WHO classification^12^.

| **Characteristics\*** | **Mild cases**<br>**(n=19)** | **Severe cases**<br>**(n=37)** | **Total cases**<br>**(n=56)** |
| --- | --- | --- | --- |
| Median age (IQR) – yr | 49 (34 - 60) | 68 (57 - 79) | 61 (48 - 77) |
| Male sex – no. (%) | 8 (42) | 23 (62) | 31 (55) |
| **COVID-19 disease severity ‡** |  |  |  |
| Mild illness– no. (%) | 12 (63) | - | 15 (27) |
| Mild pneumonia – no. (%) | 7 (37) | - | 7 (13) |
| Severe pneumonia – no. (%) | - | 21 (57) | 21 (38) |
| Mild ARDS – no. (%) | - | 4 (11) | 4 (7) |
| Moderate ARDS – no. (%) | - | 7 (19) | 7 (13) |
| Severe ARDS – no. (%) | - | 5 (14) | 5 (9) |
| **Level of care at blood sampling**<br>**timepoint** |  |  |  |
| Outpatient – no. (%) | 10 (53) | - | 10 (18) |
| Hospitalized – no. (%) | 9 (47) | 37 (100) | 46 (82) |
| **Comorbidities** |  |  |  |
| Hypertension – no. (%) | 3 (16) | 22 (59) | 25 (45) |
| Diabetes – no. (%) | 1 (5) | 10 (27) | 11 (20) |
| Heart disease – no. (%) | 1 (5) | 15 (41) | 16 (29) |
| Cerebrovascular disease – no. (%) | 1 (5) | 4 (11) | 5 (9) |
| Lung disease – no. (%) | 2 (11) | 6 (16) | 8 (14) |
| Kidney disease – no. (%) | 4 (21) | 11 (30) | 15 (27) |
| Malignancy – no. (%) | - | 4 (11) | 4 (7) |
| Immunosuppression – no. (%) | 3 (16) | 4 (11) | 7 (13) |
| * IQR denotes the interquartile range<br>‡ COVID-19 severity according to WHO guidelines<sup>12</sup> |  |  |  |

**Table 2.** Linear models for IgA and IgG serum level prediction. (**A–B**) Multiple linear model of S1 protein-specific IgA serum levels (A; logarithmized) and IgG serum levels (B; logarithmized) as a function of disease severity (mild versus severe), days since onset of symptoms, patient age, the presence of comorbidities, such as hypertension, diabetes mellitus, heart disease, cerebrovascular disease, lung disease, kidney disease, and malignancy, as well as current immunosuppressive treatment.

| SARS-CoV-2-specific IgA serum levels | Coefficient | 95% - Confidence Interval | p-value |
| --- | --- | --- | --- |
| Intercept | -0.76 | -1.89 to 0.37 | 0.18 |
| Severe disease | 1.31 | 0.60 to 2.01 | 0.0005 |
| Days | 0.05 | 0.03 to 0.07 | 0.0001 |
| Age | 0.011 | -0.01 to 0.03 | 0.37 |
| Hypertension | 0.081 | -0.79 to 0.94 | 0.85 |
| Diabetes mellitus | -0.61 | -1.44 to 0.22 | 0.15 |
| Heart disease | 0.0018 | -0.80 to 0.80 | 1.00 |
| Lung disease | -0.32 | -1.18 to 0.53 | 0.45 |
| Malignancy | -1.04 | -2.26 to 0.19 | 0.095 |
| Cerebrovascular disease | 0.98 | -0.19 to 2.16 | 0.099 |
| Kidney disease | -0.041 | -0.86 to 0.78 | 0.92 |
| Immunosuppression | -0.85 | -1.73 to 0.02 | 0.055 |

Table 2
| SARS-CoV-2-specific IgG serum levels | Coefficient | 95% - Confidence Interval | p-value |
| --- | --- | --- | --- |
| Intercept | -1.52 | -2.77 to -0.27 | 0.019 |
| Severe disease | 1.25 | 0.47 to 2.03 | 0.002 |
| Days | 0.066 | 0.04 to 0.09 | < 0.0001 |
| Age | 0.0025 | -0.02 to 0.03 | 0.85 |
| Hypertension | 0.29 | -0.66 to 1.25 | 0.54 |
| Diabetes mellitus | -0.54 | -1.47 to 0.38 | 0.24 |
| Heart disease | 0.011 | -0.88 to 0.90 | 0.98 |
| Lung disease | -0.41 | -1.37 to 0.54 | 0.39 |
| Malignancy | -0.35 | -1.71 to 1.01 | 0.60 |
| Cerebrovascular disease | -0.014 | -1.32 to 1.29 | 0.98 |
| Kidney disease | 0.32 | -0.60 to 1.23 | 0.49 |
| Immunosuppression | -1.20 | -2.17 to -0.23 | 0.016 |

**Figure 1.**
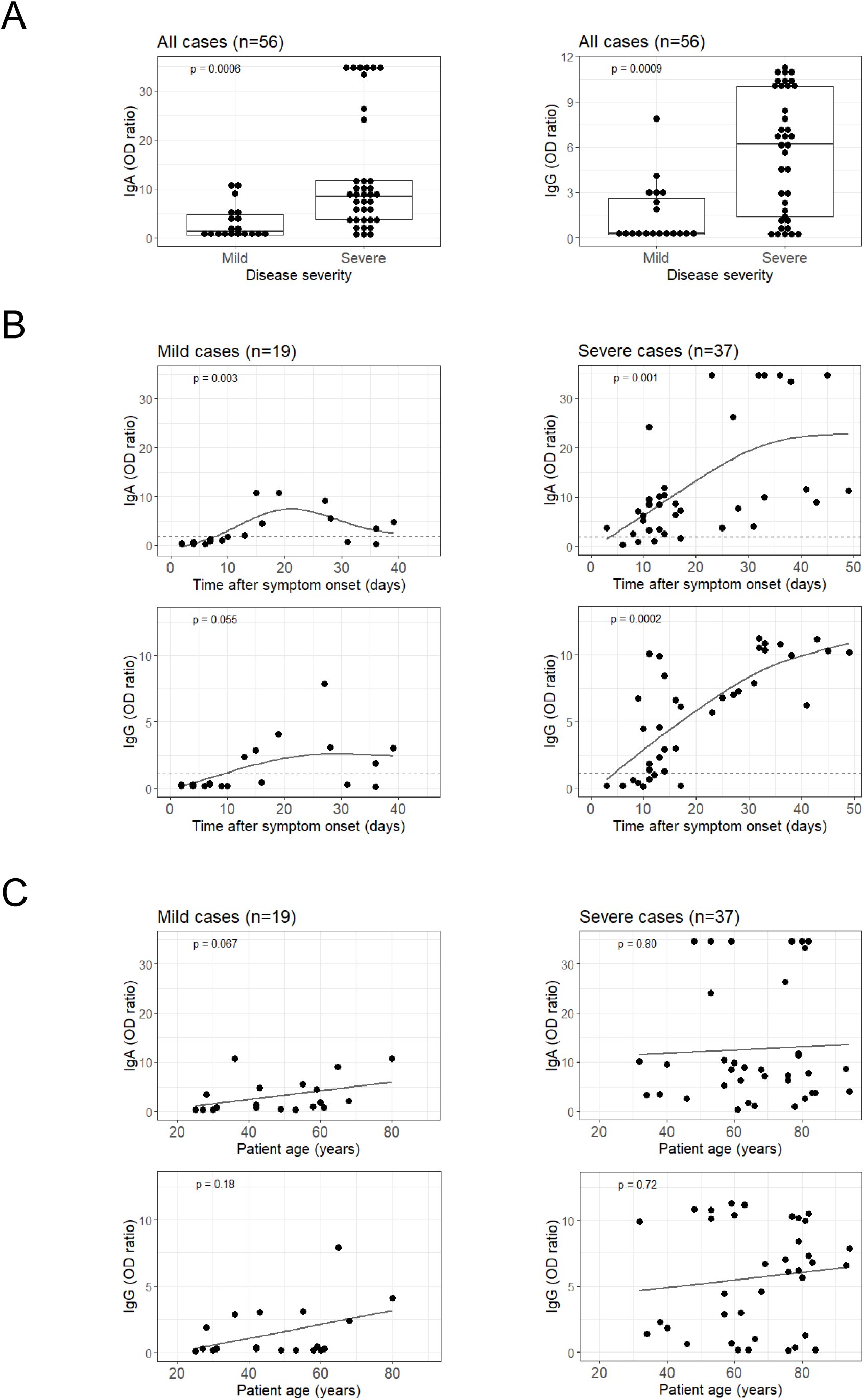
Influence of COVID-19 severity, disease duration, and patient age on SARS-CoV-2-specific serum IgA and IgG levels. **(A)** Comparison of SARS-CoV-2 spike (S) protein subunit S1-specific serum IgA and IgG titers (optical density [OD] ratio) as measured by enzyme-linked immunosorbent assay in mild (n = 19) versus severe COVID-19 cases (n = 37). Average time between reported symptom onset and sample collection was 16.4 (median 13 days) in mild and 20.9 (median 16 days) days in severe cases. **(B)** Generalized additive modeling of S1-specific IgA and IgG serum levels as a function of days between reported symptom onset and sample collection in mild (n = 19) versus severe COVID-19 cases (n = 37). Dashed lines indicate borders between positive and borderline/negative serum values of S1-specific IgA (top) and IgG (bottom). **(C)** Linear modeling of S1-specific IgA and IgG serum levels as a function of patient age in mild (n = 19) versus severe cases (n = 37). The data are shown as boxplots or scatterplots. Each dot represents an independent and unrelated donor. P-values in the between-group comparisons were calculated using Wilcoxon-test and p-values of linear and generalized additive models were computed with logarithmized IgA/IgG levels.

On average, positive S1-specific serum IgA titers became evident in mild COVID-19 cases eight days after symptom onset (Figure 1B). S1-specific serum IgA levels peaked in samples drawn at around three weeks from symptom onset, whereas in subjects tested later S1-specific serum IgA tended to be lower. As for S1-specific serum IgG concentrations, these remained negative or reached positive values in mild COVID-19 patients on average 9–10 days after symptom onset (Figure 1B).

In stark contrast to mild cases, patients suffering from severe COVID-19 showed a strong correlation of serum titers of S1-specific IgA with disease duration (p = 0.001), which was even more pronounced for serum titers of S1-specific IgG (p = 0.0002) (Figure 1B). These antibody responses became positive on average in samples obtained on day 3–4 for IgA and day 4–5 for IgG, and they appeared to be independent of patient age (p = 0.80 for IgA, and p = 0.72 for IgG), gender, and comorbidities (Figure 1B and **1C**, **Supplementary Figure S1B**, **S1C** and **S2**, and Table 2).

When grouped according to level of care into outpatient care and hospitalization, we observed in cases with mild COVID-19 that S1-specific serum IgA levels did not show any discernible pattern (Figure 2A). Contrarily, S1-specific serum IgG titers clustered according to level of care with hospitalized patients showing higher IgG titers than outpatient cases (Figure 2A), likely reflecting disease severity. Thus, we next assessed disease severity, grouping the patients according to the WHO classification criteria into mild illness without pneumonia, mild pneumonia, severe pneumonia, mild ARDS, moderate ARDS, and severe ARDS. As expected, younger patients tended to have milder disease, and older patients more severe manifestations (**Supplementary Figure S3**). There was no time-dependent pattern visible for S1-specific serum IgA, whereas S1-specific serum IgG titers showed a stronger increase over time in mild pneumonia cases versus those with mild illness (Figure 2B). Strikingly, very high levels (>25 optical density [OD] ratio) of SARS-CoV-2-specific serum IgA, but not serum IgG, correlated with severe ARDS (Figure 2B and **Supplementary Figure S4**). In a multiple linear model on all patients, there was strong evidence for an association of severe disease and days after symptom onset with an increase of S1-specific serum IgA and IgG responses. Immunosuppressive therapy was weakly associated with decreased of S1-specific serum IgG levels (Table 2).

**Figure 2.**
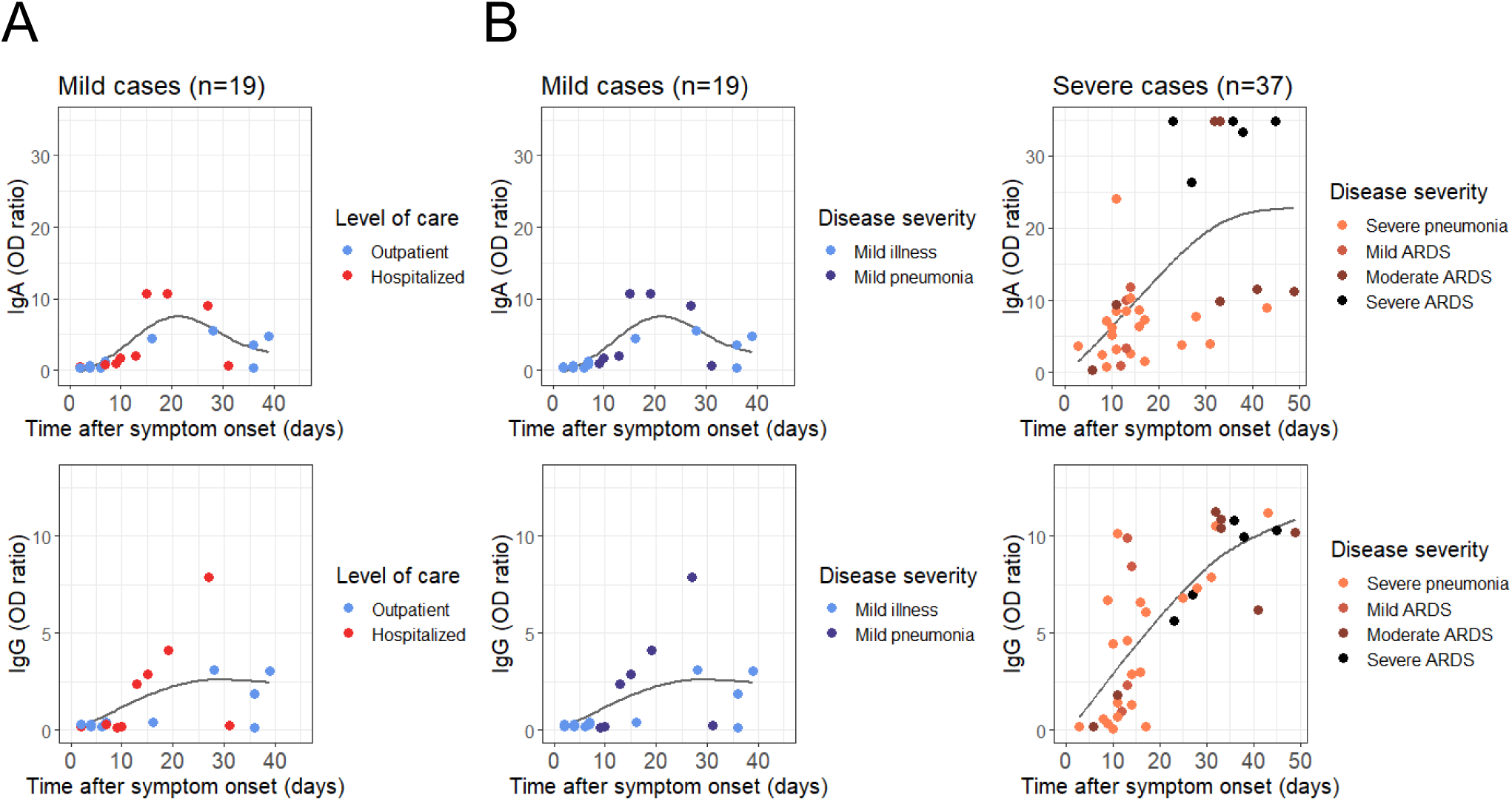
Comparison of S protein-specific serum antibodies with level of care and disease severity. **(A)** Level of patient care at the time of blood sampling, visualized in the generalized additive models of S1-specific IgA and IgG serum levels as a function of days between sampling and reported symptom onset in mild cases (n = 19). Severe cases were all hospitalized and are thus not depicted. **(B)** Disease severity at the time of blood sampling, visualized in the generalized additive models of S1-specific IgA and IgG serum levels as a function of days between sampling and reported symptom onset. Comparison of mild (n = 19) versus severe cases (n = 37).

In summary, these data show that disease severity influences S1-specific serum IgA and IgG titers, and S1-specific serum IgA of 25 OD ratio and higher might serve as a biomarker of severe ARDS. Furthermore, the data suggest that S1-specific IgA responses might occur transiently in patients with mild disease. To evaluate this latter hypothesis, we conducted a longitudinal study in two selected patients with mild COVID-19, as presented below.

### S1-specific antibody responses can be transient and delayed in mild COVID-19

We followed up two adults (a 42-year old woman and a 42-year old man, living together as a couple) with mild, RT-qPCR-confirmed SARS-CoV-2 infection. He (patient COV2-A0013) developed fatigue and cough from day 0 on, followed by fever on day 1 and dysosmia on days 9–16. She (patient COV2-A0014) showed signs of fatigue and sore throat from day 0 on, fever between days 2–5, and cough on day 3 (Figure 3A).

**Figure 3.**
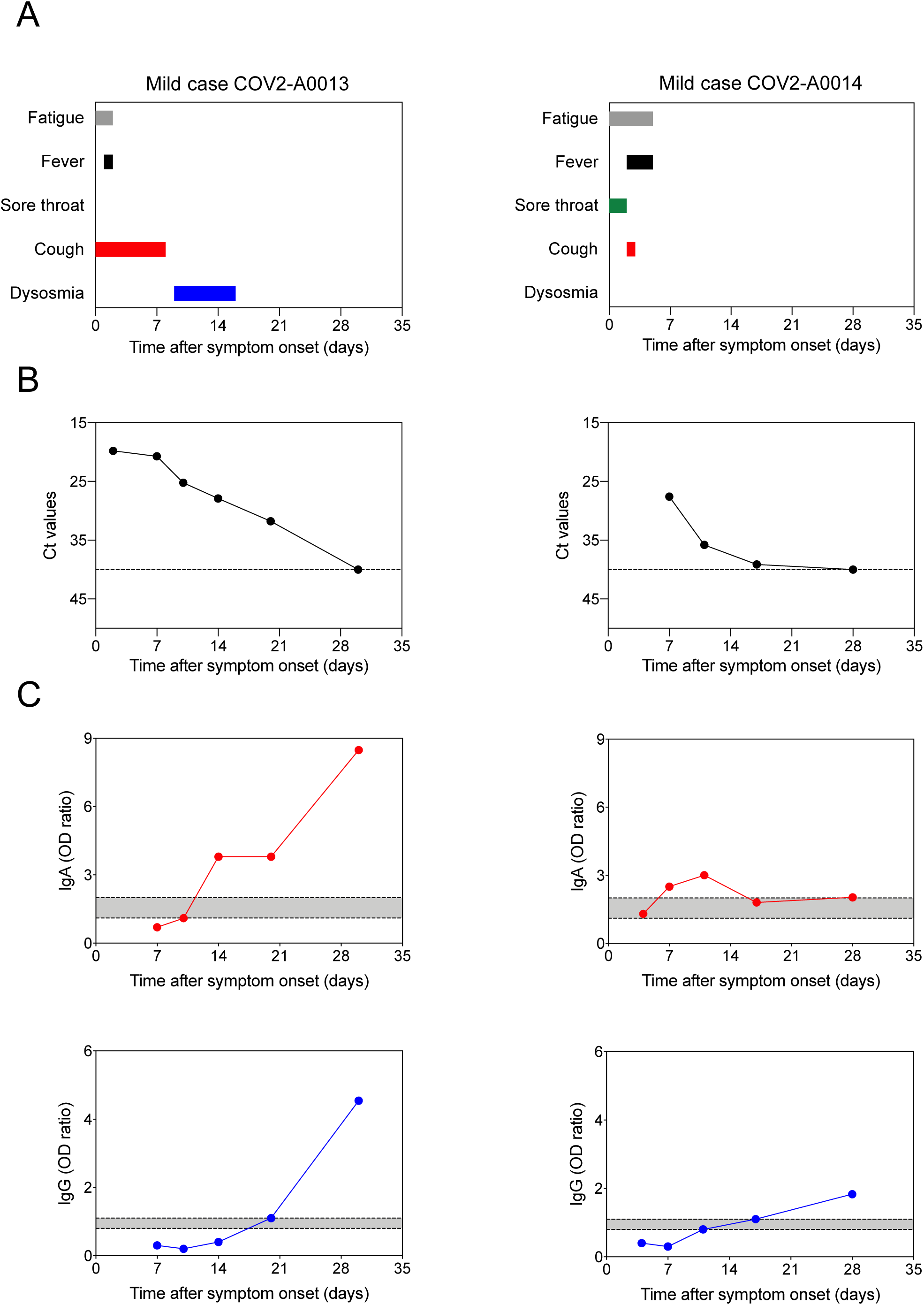
Longitudinal study of two mild cases of COVID-19. **(A)** Timeline of symptoms reported, including fatigue, fever, sore throat, cough and dysosmia, in two mild COVID-19 patients, including a 42-year old male (COV2-A0013; left panels) and a 42-year old female (COV2-A0014; right panels). **(B)** Cycle threshold (Ct) values of the SARS-CoV-2 quantitative reverse-transcriptase polymerase chain reaction (RT-qPCR) assay performed on nasopharyngeal swabs. **(C)** S1-specific IgA and IgG serum levels analyzed on different days after symptom onset. Data are shown on a longitudinal axis. Dashed lines indicate cut-offs for positive, borderline and negative serum values of S1-specific IgA (top) and IgG (bottom), with the gray shaded area highlighting the borderline values. Each dot represents an independent and unrelated measurement of the indicated patient.

The RT-qPCR cycle threshold (Ct) values at detection were low on days 1–20 for patient COV2-A0013 and day 7 for patient COV2-A0014, indicating the presence of high amounts of SARS-CoV-2 RNA in their nasal swabs (Figure 3B). On day 30 for patient COV2-A0013 and from day 17 on for patient COV2-A0014, the Ct values increased to 40 and more, thus indicating that the amount of virus RNA had dropped below the detection limit (Figure 3B).

Patient COV2-A0013 showed S1-specific serum IgA titers that were negative on day 7, rose to borderline values on day 10, became positive on day 14 at a titer of 3.8 OD ratio where they remained on day 20 and further increased to a titer of 8.5 OD ratio on day 30; his S1-specific serum IgG titers remained negative on days 7–14, became borderline positive on day 20 and clearly positive at an OD ratio of 4.5 on day 30 (Figure 3C). Conversely, S1-specific serum IgA titers in patient COV2-A0014 were borderline on day 4, became positive on days 7 and 11, followed by a drop to borderline values on days 17 and 28; her S1-specific serum IgG levels were negative on days 4– 7, became borderline on day 11 and weakly positive at an OD ratio of 1.1 on day 17 and remained weakly positive at an OD ratio of 1.8 on day 28 (Figure 3C). We compared these results to longitudinal analyses of S1-specific serum IgA and IgG values in two different situations. In asymptomatic control healthcare workers, S1-specific serum IgA and IgG remained negative throughout the period of assessment, whereas both antibodies showed a rapid increase from day 4–5 on in severe COVID-19 cases (**Supplementary Figure S5**).

Together with our aforementioned results of the 19 RT-qPCR-confirmed mild COVID-19 cases these longitudinal data in two mild COVID-19 cases demonstrate that S1-specific serum IgA production can be transient, whereas S1-specific serum IgG production occurs late, usually 17–20 days after onset of symptoms or can remain negative. Thus, collectively, these findings show that the amount of S1-specific SARS-CoV-2 IgG serum levels correlate with the severity of clinical symptoms.

### Individuals with possible SARS-CoV-2 exposure show virus-specific IgA at mucosal sites without evidence of virus-specific antibodies in serum

Having observed that in mild COVID-19 cases, S1-specific serum IgA and IgG production can be transient, delayed or even absent, we assessed serum S1-specific antibody responses in a well-defined cohort of healthcare workers possibly exposed to SARS-CoV-2 (n = 109; termed HCW cohort). These healthcare workers either did or did not have clinical symptoms suggestive of COVID-19, and they either tested negative or positive for SARS-CoV-2 in respiratory secretions by RT-qPCR. We grouped them as follows (Figure 4): (i) asymptomatic, RT-qPCR negative (Asymp/PCR-; n = 17); (ii) symptomatic, RT-qPCR negative (Symp/PCR-; n = 71); and (iii) symptomatic, RT-qPCR positive (Symp/PCR+; n = 21).

**Figure 4.**
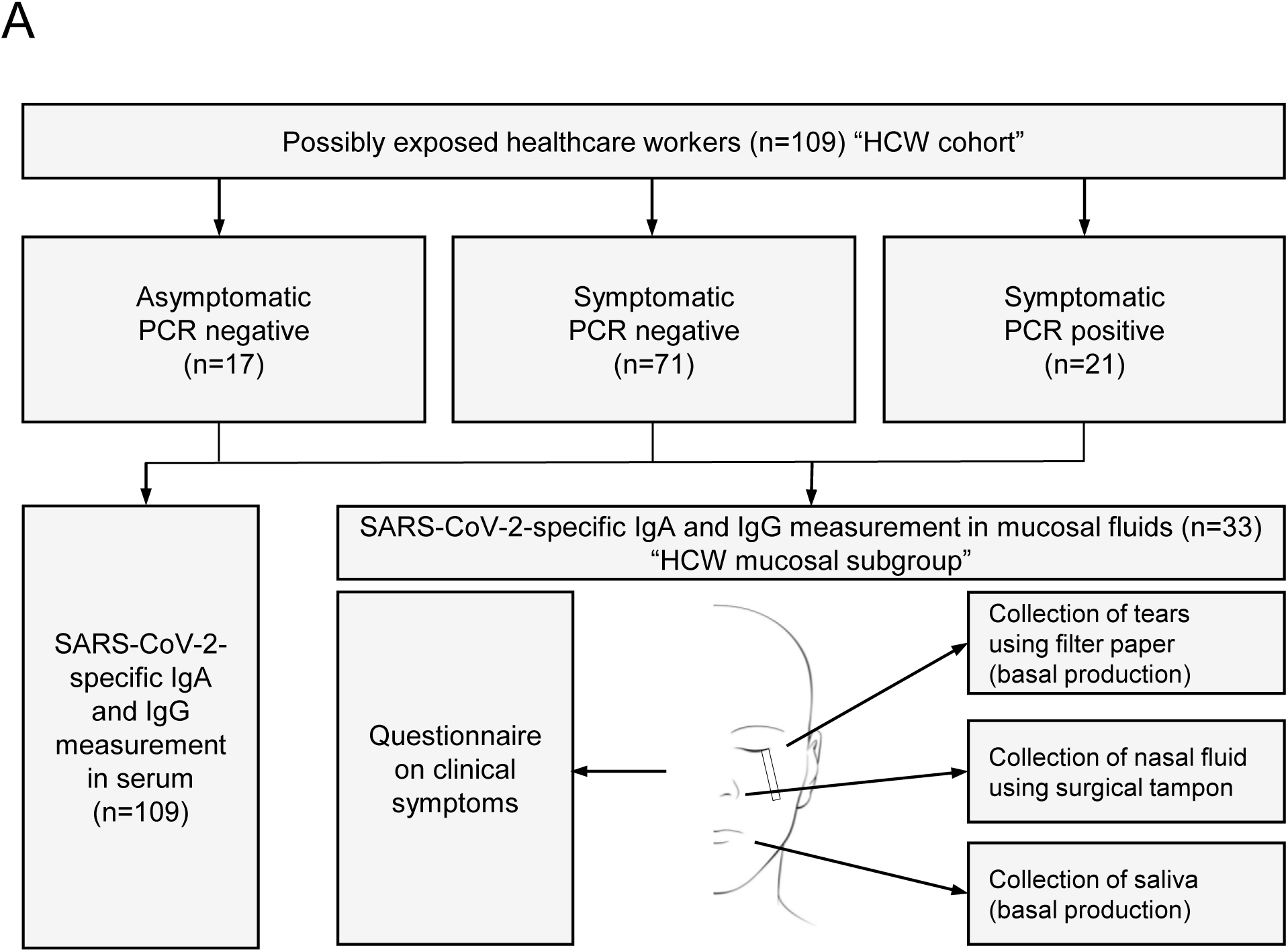
Flow chart showing characterization of a cohort of healthcare workers possibly exposed to SARS-CoV-2. (**A**) Healthcare workers (n = 109; termed HCW cohort) possibly exposed to SARS-CoV-2 were grouped as follows: (i) asymptomatic, PCR negative (Asymp/PCR–; n = 17) reporting possible exposure (see Methods) with a COVID-19 patient 11–24 days prior to sampling; (ii) symptomatic, PCR negative (Symp/PCR–; n = 71); and (iii) symptomatic, PCR positive (Symp/PCR+; n = 21). All healthcare workers were analyzed for SARS-CoV-2 S1-specific serum IgA and IgG values. In a subgroup (termed HCW mucosal subgroup), tears, nasal fluids and saliva were collected simultaneously at the time of blood draw and analyzed. Self-reported symptoms were recorded of each participant of the HCW mucosal subgroup. The 33 healthcare workers of the HCW mucosal subgroup were grouped the same way as the HCW cohort: (i) Asymp/PCR-(n = 9), tested more than 11 days after exposure; (ii) Symp/PCR-(n = 13), assessed on average 26.5 days after symptom onset; and (iii) Symp/PCR+ (n = 11), sampled on average 26.5 days after symptom onset.

The Asymp/PCR-group contained very few S1-specific serum IgA-positive and no IgG-positive subjects (Figure 5A). Conversely, there were four out of 71 (6%) participants with positive IgA and IgG values found in the Symp/PCR-group, which likely represented individuals that had had a mild SARS-CoV-2 infection (Figure 5A). As expected, the Symp/PCR+ group contained more seropositive individuals, with 8 out of 21 (38%) subjects having positive IgA and IgG titers for S1 of SARS-CoV-2 at the time of sampling (Figure 5A).

**Figure 5.**
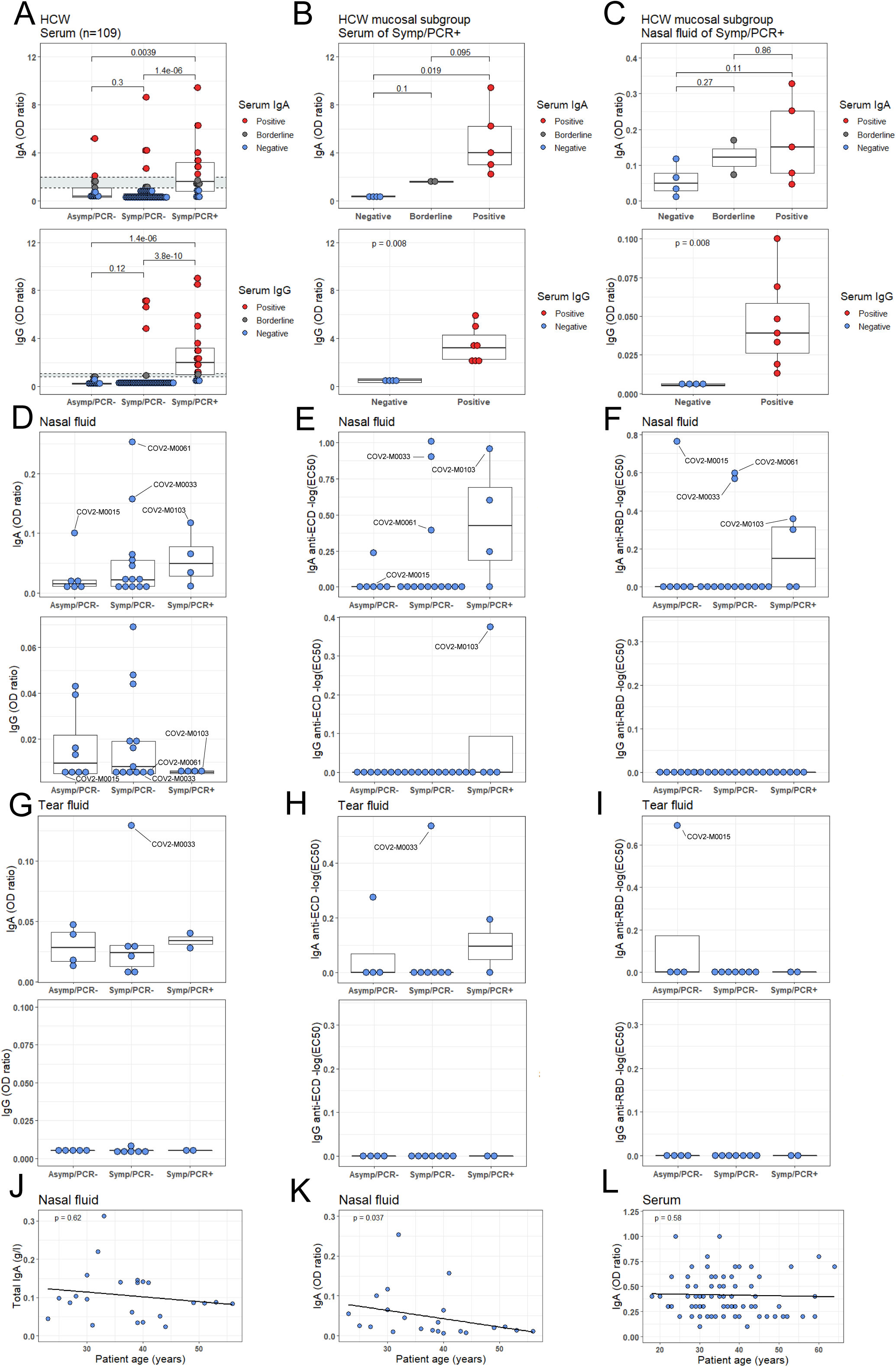
Analysis of SARS-CoV-2 S protein-specific IgA and IgG response in serum versus mucosal fluids. **(A)** S protein-specific IgA (top) and IgG (bottom)serum levels of the HCW cohort (n = 109). Dashed lines indicate borders between positive (red), borderline (gray) and negative (blue) values, with gray shaded area highlighting borderline values. **(B)** S protein-specific IgA (top) and IgG (bottom) serum levels of the Symp/PCR+ group (n = 11) in the HCW mucosal subgroup. Comparison of healthcare workers with negative, borderline, and positive values. **(C)** S protein-specific IgA (top) and IgG (bottom) nasal fluid levels of the Symp/PCR+ group (n = 11) in the HCW mucosal subgroup. Comparison of healthcare workers with negative, borderline, and positive values. (**D–F**) S protein-specific IgA (top) and IgG (bottom) levels in nasal fluid, including S1-specific IgA and IgG (D), SARS-CoV-2 S protein extracellular domain (ECD)-specific IgA and IgG (E), SARS-CoV-2 S1 protein receptor-binding domain (RBD)-specific IgA and IgG (F) of S1 protein-seronegative individuals of the HCW mucosal subgroup. Comparison of Asymp/PCR-, Symp/PCR- and Symp/PCR+ HCW is shown. Four PCR negative healthcare workers with negative S protein-specific IgA values in serum, but increased S protein-specific IgA levels in nasal fluids are labeled with their corresponding study code. (**G–I**) S protein-specific IgA (top) and IgG (bottom) levels in tear fluid, including S1-specific IgA and IgG (G), ECD-specific IgA and IgG (H), RBD-specific IgA and IgG (I) of S1 protein-seronegative individuals of the HCW mucosal subgroup. (**J–L**) Linear modeling of total polyspecific IgA in nasal fluids (J) and S1 protein-specific IgA in nasal fluids (K), and of S1 protein-specific IgA serum levels (L), all as a function of patient age in S1 protein-seronegative individuals of the HCW mucosal subgroup. Data are shown as boxplots or scatterplots and each dot represents an independent and unrelated donor. P-values of linear additive models were computed with logarithmized IgA/IgG levels. P-values of the between-group differencies were calculated using Wilcoxon-test.

To investigate S1-specific IgA and IgG levels at mucosal sites, we analyzed tears, nasal fluids, and saliva in a subset of the HCW cohort, termed HCW mucosal subgroup (Figure 4). This subgroup also recorded self-reported clinical symptoms related to their possible SARS-CoV-2 exposure (Table 3). When assessing the Symp/PCR+ members of the HCW mucosal subgroup, a clear correlation was evident between positivity of S1-specific IgA and IgG in serum (Figure 5B) with the corresponding values in nasal secretions (Figure 5C). Thus, for S1-specific IgG, Symp/PCR+ members with positive serum titers also showed elevated levels of S1-specific IgG in nasal secretions (Figure 5B and 5C), possibly indicating transfer of S1-specific IgG from serum to the nasal mucosa. Conversely, the relationship of serum versus nasal fluid in Symp/PCR+ members was more variable for S1-specific IgA (Figure 5B and 5C).

**Table 3.** Demographic and clinical characterstics of the healthcare worker cohort. **(A)** Healthcare worker cohort (HCW cohort) included in the S protein-specific IgA and IgG serology study. **(B)** Healthcare worker subgroup (HCW mucosal subgroup) assessed in the S protein-specific IgA and IgG mucosal fluid study.

| **HCW cohort** |  |  |  |  |
| --- | --- | --- | --- | --- |
| **Characteristics \*** | **Asymptomatic/**<br>**PCR- (n=17)** | **Symptomatic/**<br>**PCR- (n=71)** | **Symptomatic/**<br>**PCR+ (n=21)** | **Total**<br>**(n=109)** |
| Median age (IQR) – yr | 39 (34 - 44) | 36 (30 - 41) | 38 (30 - 48) | 36 (30 - 43) |
| Male sex – no. (%) | 6 (35) | 16 (23) | 3 (14) | 25 (23) |
| * IQR denotes the interquartile range |  |  |  |  |

Table 3
| **HCW mucosal subgroup** |  |  |  |  |
| --- | --- | --- | --- | --- |
| **Characteristics \*** | **Asymptomatic/**<br>**PCR- (n=9)** | **Symptomatic/**<br>**PCR- (n=13)** | **Symptomatic/**<br>**PCR+ (n=11)** | **Total**<br>**(n=33)** |
| Median age (IQR) – yr | 38 (36 - 44) | 40 (32 - 49) | 38 (30 - 42) | 39 (31 - 43) |
| Male sex – no. (%) | 4 (44) | 6 (46) | 3 (27) | 13 (41) |
| **Reported symptoms** |  |  |  |  |
| Fatigue – no. (%) | - | 6 (46) | 7 (64) | 13 (39) |
| Body temperature<br>> 38.0°C – no. (%) | - | 4 (31) | 1 (9) | 5 (15) |
| Feeling feverish – no. (%) | - | 6 (46) | 4 (36) | 10 (30) |
| Chills – no. (%) | - | 1 (8) | 2 (18) | 3 (9) |
| Shivering – no. (%) | - | 3 (23) | 4 (36) | 7 (21) |
| Body aches – no. (%) | - | 8 (62) | 8 (73) | 16 (48) |
| Back pain – no. (%) | - | 5 (38) | 4 (36) | 9 (27) |
| Cough – no. (%) | - | 5 (38) | 6 (55) | 11 (33) |
| Dyspnea – no. (%) | - | 2 (15) | 4 (36) | 6 (18) |
| Pleuritis – no. (%) | - | 3 (23) | 4 (36) | 7 (21) |
| Sore throat – no. (%) | - | 11 (85) | 6 (55) | 17 (52) |
| Coryza – no. (%) | - | 7 (54) | 6 (55) | 13 (39) |
| Hoarseness – no. (%) | - | 5 (46) | 4 (36) | 9 (27) |
| Anosmia/Dysosmia – no. (%) | - | 2 (15) | 8 (73) | 10 (30) |
| Diarrhea – no. (%) | - | 5 (38) | 2 (18) | 7 (21) |
| Nausea – no. (%) | - | 3 (23) | 3 (27) | 6 (18) |
| Conjunctivitis – no. (%) | - | 2 (15) | - | 2 (6) |
| * IQR denotes the interquartile range |  |  |  |  |

To further investigate these findings, we adapted and used our two SARS-CoV-2 S protein-specific immunoassays (**Supplementary Figure S6** and **S7**) to assess the subjects in the HCW mucosal subgroup that tested negative for SARS-CoV-2-specific IgA or IgG in serum. Firstly, we ruled out an influence by sampling time since symptom onset or total amount of detectable polyspecific IgA and IgG in our samples. Time of sampling since symptom onset was 26.5 days for both Symp/PCR- and Symp/PCR+ groups of the HCW mucosal subgroup, whereas the Asymp/PCR-group was tested more than 11 days after exposure. Total polyspecific IgA and IgG levels were comparable in serum samples as well as tear, nasal fluid and saliva samples of all three groups of participants (**Supplementary Figure S8**). Notably, whereas total polyspecific IgG levels were measurable in nasal fluids, they were very low in tear fluid and saliva (**Supplementary Figure S8**). Analyzing S protein-specific IgA and IgG in our mucosal samples, we observed a reliable correlation between our two immunoassays for serum IgA and IgG, tear fluid IgA, and nasal fluid IgA, whereas the other measurements were not consistent and were thus not considered for our conclusions (**Supplementary Figure S7**).

Interestingly, we were able to detect S protein-specific IgA in the mucosal samples of several subjects in the absence of seropositivity. Analyzing individual participants, we found that subjects COV2-M0033, COV2-M0061, and COV2-M0103 showed high S1-specific, ECD-specific and RBD-specific IgA values in their nasal fluids, whereas total IgA values were average in nasal fluids of these individuals (Figure 5D–5F and **Supplementary Figure S8**). Moreover, the nasal fluid of subject COV2-M0015 contained high S1-specific and RBD-specific IgA values, in the presence of average total IgA values (Figure 5D–5F and **Supplementary Figure S8**). When measuring their tear fluids, subjects COV2-M0015 and COV2-M0033 presented with high S1-specific, ECD-specific or RBD-specific IgA values (Figure 5G–5I). Additionally, a few other individuals also had detectable S protein-specific IgG in their nasal fluids despite being IgG seronegative (Figure 5D–5F).

In contrast to total IgA values, assessing S protein-specific IgA values in nasal fluid versus age in seronegative healthcare workers, we found an inverse correlation (p = 0.037) (Figure 5J and 5K). The same analysis with S protein-specific IgA titers in serum versus age, however, did not reveal a correlation (p = 0.58) (Figure 5L).

Collectively, these data suggest that mucosal SARS-CoV-2 S protein-specific IgG antibody levels are related to systemic titers of these antibodies. Interestingly, in 15–20% of S protein-seronegative individuals, we were able to detect S protein-specific IgA antibodies at several mucosal sites. Furthermore, mucosal S protein-specific IgA levels inversely correlated with patient age, suggesting increased mucosal antibody responses in younger SARS-CoV-2-exposed individuals.

## Discussion

In severe cases of COVID-19, we found that SARS-CoV-2 S protein-specific serum IgA and IgG titers became positive in samples obtained on average 3–5 days after symptom onset, which is in agreement with earlier publications.^14^ These antibody responses showed a strong correlation with disease duration, but they were independent of patient age, gender, and pre-existing comorbidities. Strikingly, very high serum titers of S protein-specific IgA, but not IgG, correlated with severe ARDS, thus warranting further studies evaluating the role of IgA in SARS-CoV-2-associated severe ARDS.

Conversely, in mild cases of SARS-CoV-2 infection, S protein-specific serum IgA production was transient, delayed or even absent, accompanied by an S protein-specific serum IgG response that occurred late or remained negative. Interestingly, however, we found evidence of S protein-specific IgA and IgG at mucosal sites of individuals with mild COVID-19. There, mucosal S protein-specific IgG levels appeared to mirror the systemic, i.e. serum, titers of these antibodies. Mucosal S protein-specific IgA levels, however, were even detectable at several mucosal sites of about 15–20% S protein-seronegative individuals. Interestingly, mucosal S protein-specific IgA levels correlated inversely with patient age.

We think these findings suggest a model where the extent and duration of SARS-CoV-2-related clinical symptoms, which likely correlates with virus replication, dictates the level of virus-specific humoral immunity. This hypothesis is consistent with previous publications demonstrating that the magnitude of the humoral response toward SARS-CoV-2 is dependent on the duration and magnitude of viral antigen exposure.^26,27^ Low antigen exposure will elicit mucosal IgA-mediated responses, which can be accompanied by systemic IgA production; however, systemic virus-specific IgA responses can also be absent, transient or delayed. This type of “mucosal IgA” antibody response seemed to be particularly prevalent in younger individuals with mild SARS-CoV-2 infection without evidence of pneumonia. These projected longitudinal relationships from cross-sectional evaluations need confirmation in longitudinal studies. Notably, of our two longitudinal subjects, patient COV2-A0014 showed milder and shorter clinical symptoms and more rapid virus clearance, which was associated with transient S protein-specific IgA and delayed IgG production, but high levels of S protein-specific IgA in her nasal fluid (**Supplementary Figure S9**).

These data might be a reflection of increased mucosal immunity in the young or decreased mucosal immunity in the old. Along these lines, previous data on coronavirus seroprevalence of HKU1-specific IgG showed an absence of systemic HKU1-specific antibodies in individuals younger than 20 years of age, with increasing seroprevalence with increasing age.^28^ Extrapolating this model to comprise also children and infants, it is conceivable that children and infants have primed mucosal innate and IgA antibody responses due to their frequent upper respiratory tract infections and, therefore, respond preferentially in this manner to SARS-CoV-2 infection. This hypothesis might also explain why children rarely present with symptomatic SARS-CoV-2 infection. Looking at the other end of the age sprectrum, previous studies have shown that the kinetics and strength of antiviral immune responses, including T cell activation and proliferation, becomes slower with increasing age.^29,30^ The elucidation of these questions and the confirmation of our findings will require larger studies. However, due to the transient nature of S protein-specific antibody responses in oligosymptomatic patients, large scale measurement of systemic SARS-CoV-2-specific IgA and IgG levels in asymptomatic patients will reveal limited epidemiological information. In addition to serum, mucosal measurements of SARS-CoV-2-specific IgA should be considered.

With increased SARS-CoV-2-related clinical symptoms, and hence antigen exposure, we observed a “systemic IgA and IgG” type of antibody response, characterized by S protein-specific IgA that may be transient or delayed and the presence of S protein-specific IgG. With even further increasing clinical severity, we found high to very high serum IgA and high IgG responses in severe cases and ARDS. Thus, our findings suggest four grades of antibody responses dependent on COVID-19 severity with (i) oligosymptomatic disease and mucosal antibody responses in the absence of systemic antibody production, (ii) mild-to-moderate disease and transient or delayed systemic IgA and IgG production, (iii) severe cases with high serum IgA and high IgG responses, and (iv) very severe cases, including severe ARDS, with very high serum IgA and high IgG titers.

Whether these S protein-specific antibody responses confer immunity to a secondary infection with SARS-CoV-2 is a matter of intense debate. Previous publications indicated that S protein-specific serum IgG antibodies correlated with virus neutralization in vitro,^14,15^ although some publications questioned the efficacy of neutralization by these antibody responses.^20^ Based on correlative data from the SARS outbreak and preclinical SARS infection models,^31^ there has been discussion on a contribution of the humoral immune response to immune pathology,^32,33^ potentially by augmenting the proinflammatory monocyte response in the lungs. However, trials with convalescent serum treatments have shown promising results during the current COVID-19 pandemic and also in SARS.^34^ Another caveat relates to the durability of protective humoral immunity. Whether S protein-specific mucosal IgA responses confer immunity to a secondary infection with SARS-CoV-2 remains to be seen. We are currently assessing S protein-specific mucosal IgA antibodies in virus neutralization assays as well as following-up our patient cohort longitudinally to address these issues.

## Acknowledgments

We thank Alessandra Guaita, Jennifer Jörger, Mitchell Levesque, Hugo Sax, Urs Steiner, and the members of the Boyman laboratory for helpful discussions. We are also grateful to those healthcare workers at USZ who helped with sampling and recruitment of the HCW cohort.

## Author contributions

CC and JN contributed to study design, patient recuitment, data collection, data analysis, data interpretation, and writing of the manuscript. YZ and AW contributed to patient recuitment, clinical management, data collection, data analysis, and data interpretation. AV, JS, SA, ME, PPB, EDC, AA, and EP-M contributed to experiments, data collection, data analysis, data interpretation. MER, SH, EB, AR, MS-H, LCH, ASZ, and DJS contributed to patient recuitment and clinical management. UH contributed to data analysis and data interpretation, particularly statistcal analysis. SKR contributed to study design, patient recuitment, data collection, data analysis, and data interpretation. OB contributed to study conception and design, data analysis, data interpretation, and wrote the manuscript. All authors reviewed and approved the final version of the manuscript.

## Funding

This work was funded by Swiss Academy of Medical Sciences fellowships (#323530-191220 to CC; #323530-191230 to YZ; #323530-177975 to SA), the Young Talents in Clinical Research Fellowship by the Swiss Academy of Medical Sciences and Bangerter Foundation (YTCR 32/18 to MR), the Swiss National Science Foundation (#310030-172978 to OB), the Clinical Research Priority Program of the University of Zurich for the CRPP CYTIMM-Z (to OB), and a grant of the Innovation Fund of the University Hospital Zurich (to OB).

## Competing interests

The authors declare no competing financial interests related to this work.

## Notes

### Competing Interest Statement

The authors have declared no competing interest.

