## Supplementary Material for "Systemic and mucosal antibody secretion specific to SARS-CoV-2 during mild versus severe COVID-19"

### Supplementary Figure legends

**Supplementary Figure S1. Comparison of S protein-specific serum IgA and IgG values with sampling timepoint, disease severity, patient age, and gender.**

(**A**) Comparison of the days between reported symptom onset and sample collection in mild (n = 19) versus severe COVID-19 cases (n = 37).

(**B**) Visualization of age distribution in the generalized additive models of S1 protein-specific IgA and IgG serum levels as a function of days between reported symptom onset and sample collection. Comparison of mild (n = 19) versus severe cases (n = 37).

(**C–D**) Comparison of S1 protein-specific serum IgA (C) and IgG (D) titers in male (n = 31) versus female (n = 25) COVID-19 patients. Average days between reported symptom onset and sample collection were 20.2 (median 15) in male and 18.4 (median 14) days in female patients.

**Supplementary Figure S2. Comparison of S protein-specific serum IgA and IgG values with patient comorbidities.**

(**A**–**D**) Visualization of reported patient comorbidities (hypertension, diabetes mellitus, heart disease, cerebrovascular disease, lung disease, kidney disease, and malignancy) and current immunosuppressive treatment in the generalized additive models of S1 protein-specific IgA (A and C) and IgG (B and D) serum levels as a function of days between reported symptom onset and sample collection in mild (n = 19; A and B) and severe cases (n = 37; C and D).

**Supplementary Figure S3. Distribution of disease severity and age of COVID-19 patients.**

Comparison of COVID-19 patients' (n = 56) age distribution in relation to their COVID-19 severity at the time of sample collection, ranging from mild COVID-19 to severe ARDS, as defined by the WHO classification criteria[12]. P-value was computed using Kruskal-Wallis test.

**Supplementary Figure S4. Comparison of S protein-specific serum IgA and IgG values and symptom severity of COVID-19 patients.**

**(A–B**) Comparison of S1 protein-specific serum IgA (A) and IgG (B) titers with disease severity in our COVID-19 patient cohort (n = 56), ranging from mild COVID-19 to severe ARDS, as defined by the WHO classification criteria[12]. Data are shown as boxplots. Each dot represents an independent and unrelated donor. The significance of the between-group differencies was explored by using Kruskal-Wallis test.

**Supplementary Figure S5. Longitudinal measurement of S protein-specific serum IgA and IgG values in asymptomatic controls and severe cases of COVID-19.**

(**A–B**) S1 protein-specific serum IgA (top) and IgG (bottom) titers in asymptomatic donors (A; n = 4) and severe cases of COVID-19 (B; n = 3). The connected dots represent sequential measurements of the same individual.

**Supplementary Figure S6. Titration of nasal fluids to detect S protein-specific IgA and IgG.**

Measurement of S1 protein-specific IgA (top) and IgG (bottom) using different dilutions of nasal fluids of a subset of the HCW mucosal subgroup (n = 15).

**Supplementary Figure S7. Comparison of immunoassays to measure S protein-specific IgA and IgG in samples of serum, tears, nasal fluid, and saliva.**

Comparison of optical density (OD) ratios of IgA (top) and IgG (bottom) obtained with a commercial enzyme-linked immunosorbent assay specific for the S1 protein of SARS-CoV-2 (x-axes) and the inflection point of the sigmoidal curve (-log(EC50); y-axes), the latter determined by measuring IgA (top) and IgG (bottom) against SARS-CoV-2 S protein extracellular domain (ECD) and SARS-CoV-2 S1 protein receptor-binding domain (RBD) in serial dilutions using an in-house immunoassay (see Methods). S protein-specific IgA (top) and IgG (bottom) were measured in serum, tear fluid, nasal fluid, and saliva of the healthcare worker mucosal subgroup (HCW mucosal subgroup). Data are shown as scatterplots. Each dot represents an independent and unrelated donor.

**Supplementary Figure S8. Analysis of total IgA and IgG serum levels in HCW mucosal subgroup.**

Total polyspecific (i.e. of undefined specificity) IgA (top) and IgG (bottom) levels in serum, tear fluid, nasal fluid, and saliva were assessed in individuals of the HCW mucosal subgroup that tested negative for S1 protein-specific serum IgA (top) and IgG (bottom).

Comparison of Asymp/PCR-, Symp/PCR- and Symp/PCR+ HCW is shown. Four PCR negative healthcare workers with negative S protein-specific IgA values in serum, but increased S protein-specific IgA levels in nasal fluids are labeled with their corresponding study code. The significance of the between-group differencies was explored by using Wilcoxon test.

**Supplementary Figure S9. Mucosal S protein-specific IgA and IgG in two mild COVID-19 cases.**

Shown are S1 protein-specific IgA and IgG titers in the nasal fluids of patients COV2-A0013 and COV2-A0014 (see Figure 3) on day 25.

S1

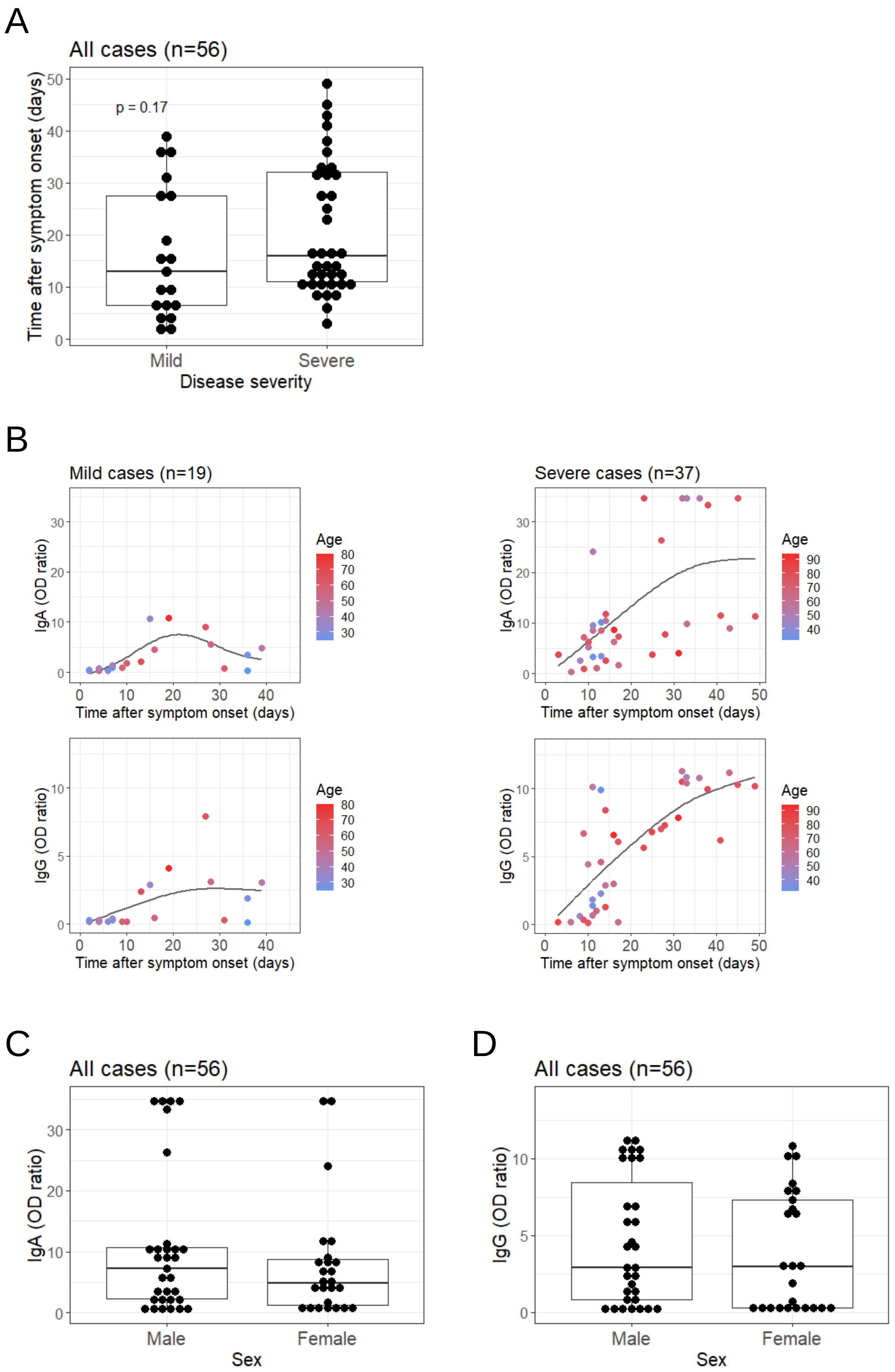


Supplementary

S2

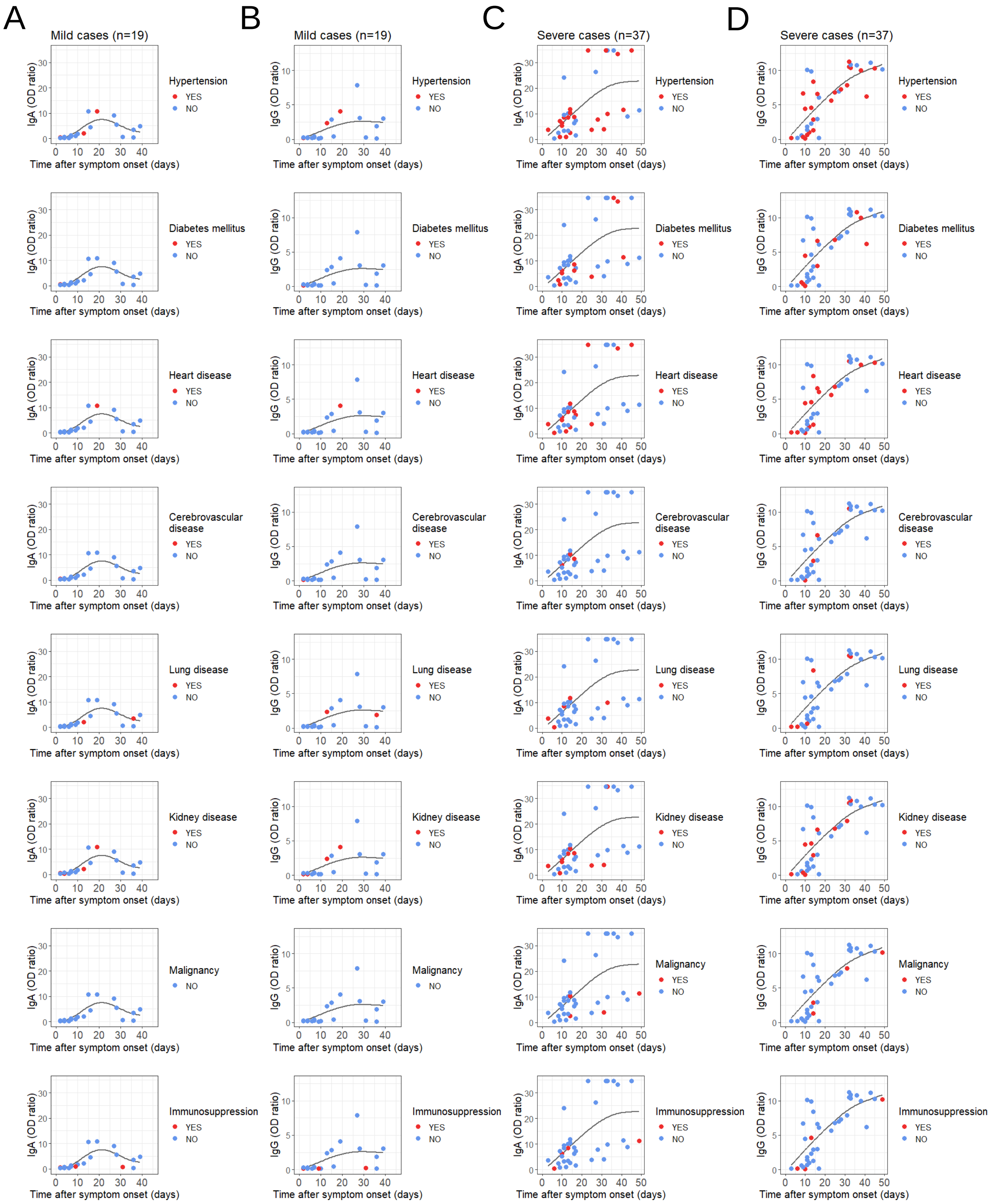


Supplementary

S3

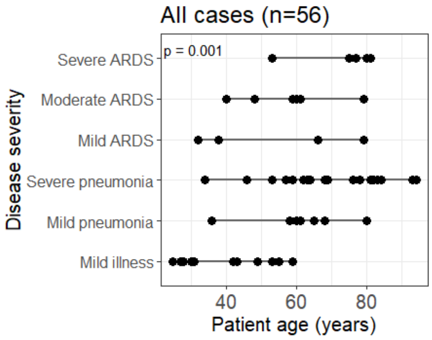
All cases (n=56)
p = 0.001
Severe ARDS
Moderate ARDS
Mild ARDS
Severe pneumonia
Mild pneumonia
Mild illness
Disease severity
40
60
80
Patient age (years)

S4

A

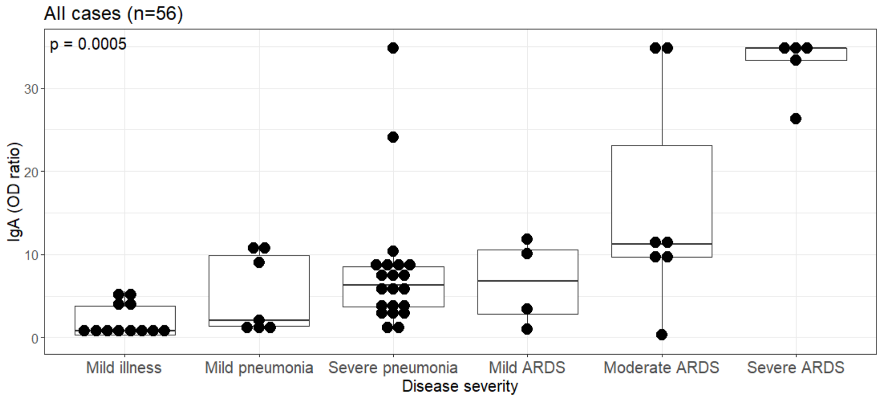
All cases (n=56)
p = 0.0005
IgA (OD ratio)
0
10
20
30
Mild illness
Mild pneumonia
Severe pneumonia
Mild ARDS
Moderate ARDS
Severe ARDS
Disease severity

B

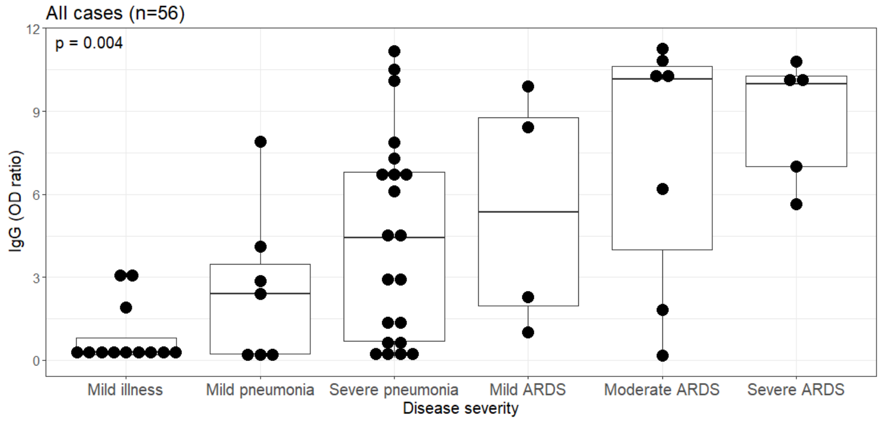
All cases (n=56)
p = 0.004
IgG (OD ratio)
0
3
6
9
12
Mild illness
Mild pneumonia
Severe pneumonia
Mild ARDS
Moderate ARDS
Severe ARDS
Disease severity

S5

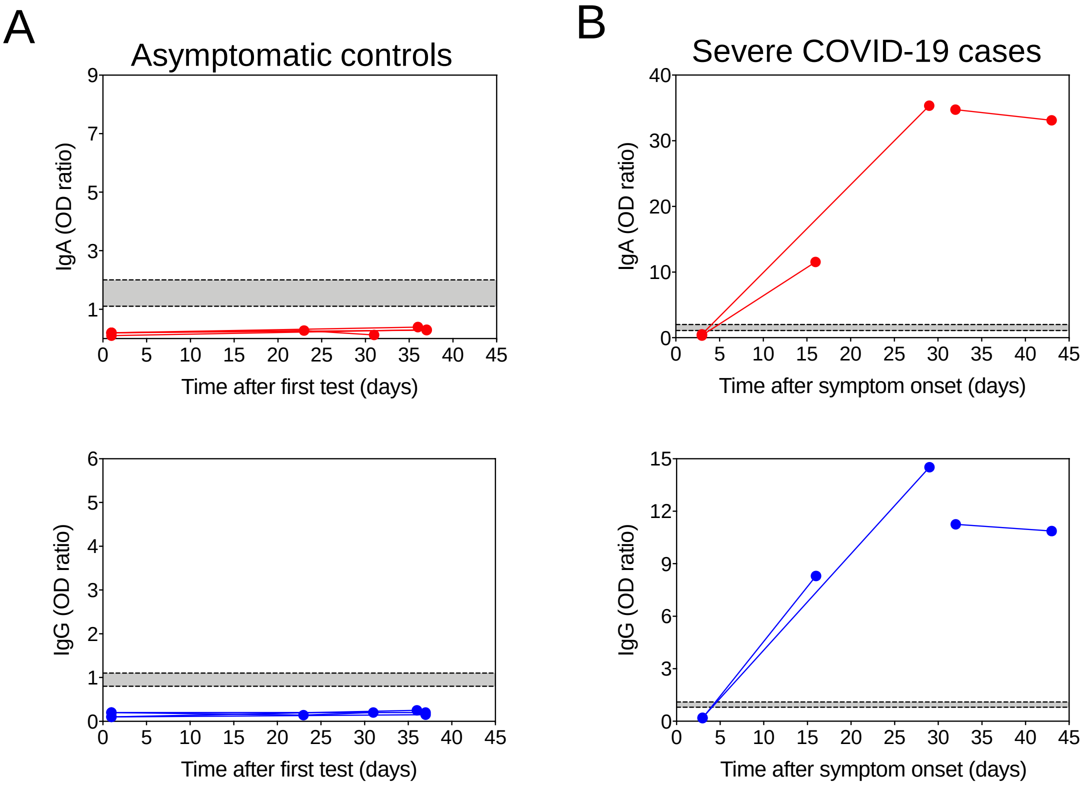
A
Asymptomatic controls
IgA (OD ratio)
Time after first test (days)
IgG (OD ratio)
Time after first test (days)
B
Severe COVID-19 cases
IgA (OD ratio)
Time after symptom onset (days)
IgG (OD ratio)
Time after symptom onset (days)

S6

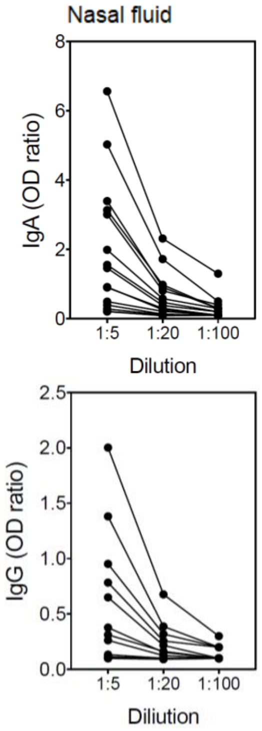
Nasal fluid
IgA (OD ratio)
1:5
1:20
1:100
Dilution
IgG (OD ratio)
1:5
1:20
1:100
Diliution

S7

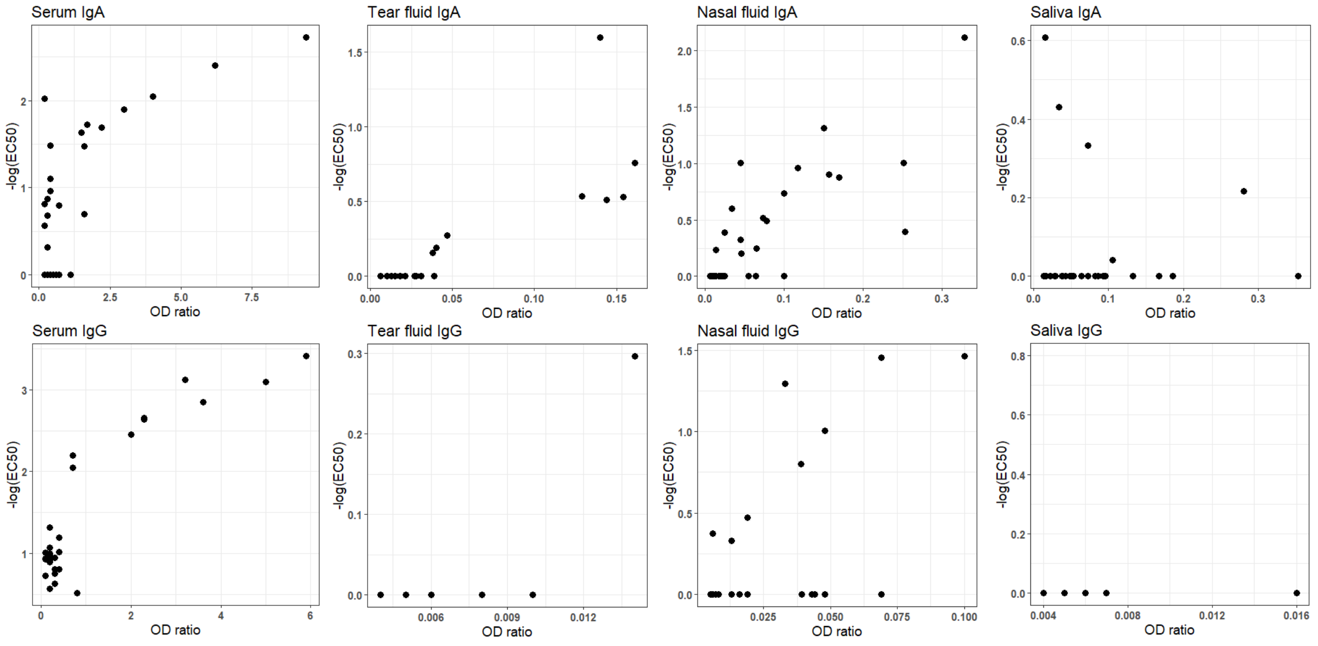


S8

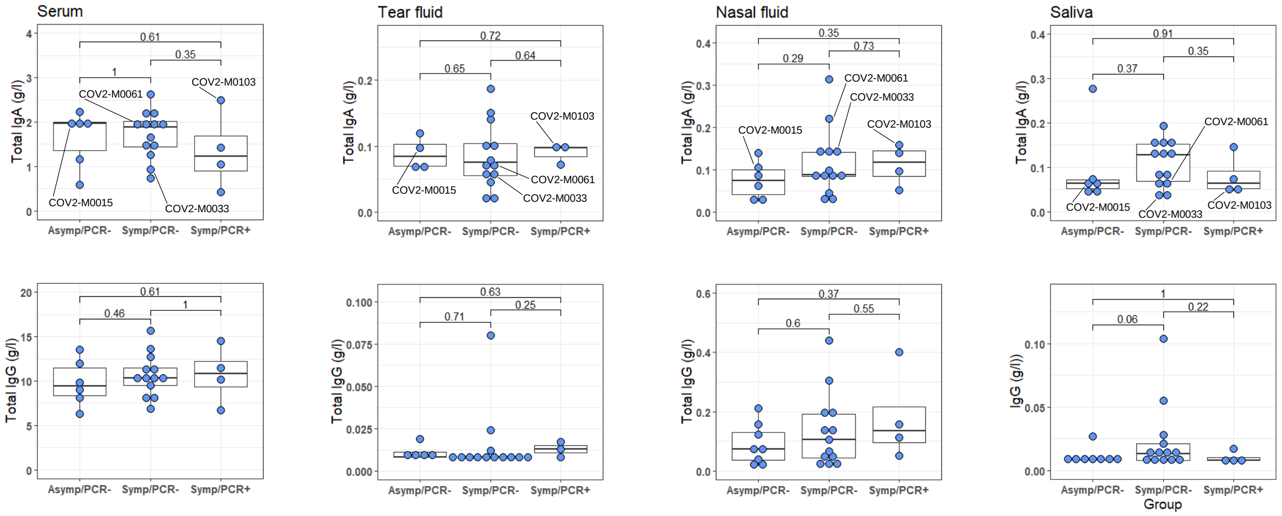


S9

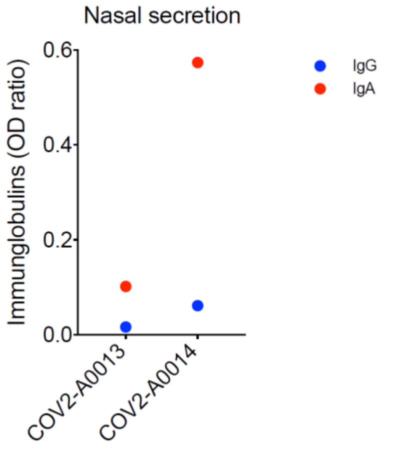


Supplementary
